# Heroin self-administration and extinction increases prelimbic cortical astroglia-synapse proximity and alters dendritic spine morphometrics that are reversed by N-acetylcysteine

**DOI:** 10.1101/2020.05.21.108530

**Authors:** BM Siemsen, AR Denton, KN Hooker, EA Carpenter, ME Prescott, AG Brock, MN Leath, JA McFaddin, JF McGinty, MD Scofield

## Abstract

Clinical and preclinical studies indicate that adaptations in corticostriatal neurotransmission significantly contribute to heroin relapse vulnerability. In animal models, heroin self-administration and extinction produce cellular adaptations in both neurons and astrocytes within the nucleus accumbens (NA) core that are required for cue-induced heroin seeking. Specifically, decreased glutamate clearance and reduced association of perisynaptic astrocytic processes with NAcore synapses allow glutamate release from prelimbic (PrL) cortical terminals to engage synaptic and structural plasticity in NAcore medium spiny neurons. Normalizing astroglial glutamate homeostasis with drugs like the antioxidant N-acetylcysteine (NAC) prevents cue-induced heroin seeking. Surprisingly, little is known about heroin-induced alterations in astrocytes or pyramidal neurons projecting to the NAcore in the PrL cortex (PrL-NAcore). Here, we observed increased complexity of the glial fibrillary acidic protein (GFAP) cytoskeletal arbor and increased association of the astroglial plasma membrane with synaptic markers following heroin SA and extinction training in the PrL cortex. Repeated treatment with NAC during extinction reversed both the enhanced astroglial complexity and synaptic association. In PrL-NAcore neurons, heroin SA and extinction decreased apical tuft dendritic spine density and enlarged dendritic spine head diameter in male Sprague-Dawley rats. Repeated NAC treatment during extinction prevented decreases in spine density but not dendritic spine head expansion. Moreover, heroin SA and extinction increased co-registry of the GluA1 subunit of AMPA receptors in both the dendrite shaft and spine heads of PrL-NAcore neurons. Interestingly, accumulation of GluA1 immunoreactivity in spine heads was further potentiated by NAC treatment during extinction. Taken together, our data reveal circuit-level adaptations in cortical dendritic spine morphology potentially linked to heroin-induced alterations in astrocyte complexity and association at synapses. Additionally, these data demonstrate, for the first time, that NAC reverses PrL cortical heroin SA and extinction-induced adaptations in both astrocytes and corticostriatal neurons.

## Introduction

Relapse vulnerability remains a major clinical obstacle in treating opioid use disorders, with relapse to drug seeking typically brought on by craving evoked by exposure to opioid-associated contexts or cues [1]. In humans, elevated activity in the corticostriatal circuit is an established biomarker of cue-evoked craving for heroin [2] and heroin-mediated dysfunction in corticostriatal circuity is a central component of the construct validity of operant animal models of heroin self-administration (SA) and relapse [3]. A general consensus from rodent models is that dysfunctional prefrontal cortical regulation of ventral striatal plasticity occurs as a result of chronic opioid exposure, allowing opioid-associated cues and contexts to drive drug seeking even after prolonged periods of abstinence [4].

The nucleus accumbens core (NAcore) is required for motivationally-relevant external stimuli to evoke motor programs linked to seeking reinforcers or rewards [5]. It has been repeatedly demonstrated that heroin SA in rodents alters glutamatergic plasticity in the NAcore in a manner that is mechanistically linked to relapse [6, 7]. For example, following extinction of heroin [8] or cocaine [9] SA, an inability to induce and maintain LTP and LTD in the NAcore is observed following stimulation of the prelimbic (PrL) cortex *in vivo*. These data demonstrate drug-mediated dysfunctional plasticity at PrL-NAcore synapses. Akin to cocaine [10], heroin SA and extinction decreases the glutamate clearance in the NAcore due to downregulation of the high-affinity astroglial glutamate transporter, GLT-1 [11] and concomitant reductions in the proximity of GLT-1-containing perisynaptic astrocyte processes (PAPs) to NAcore synapses [12]. Interestingly, decreased astrocytic association with NAcore synapses is also observed following extinction of methamphetamine or cocaine SA [13, 14]. Together these data suggest that structural and cellular astrocyte dysfunction in the NAcore is a central aspect of the pathophysiology underlying susceptibility to relapse [15]. As such, reduced GLT-1 expression coupled with decreased proximity of astrocyte PAPs to synapses likely synergistically facilitate glutamate spillover to extrasynaptic sites following synaptic release. Consistent with dysfunctional plasticity at PrL-NAcore synapses, cue- and heroin-primed seeking promotes sustained elevation of glutamate levels in the NAcore specifically due to release from PrL cortical afferents [16]. This elevated glutamate release and spillover facilitates LTP-like increases in synaptic strength, including an upregulation of NR2B-containing NMDA receptors and enlargement of dendritic spine heads, on NAcore medium spiny neurons (MSNs). Accordingly, it has been shown that normalizing drug-induced alterations in PrL-NAcore synaptic transmission, mediated in large part by astrocytic dysfunction, serves as a potent means for decreasing relapse vulnerability [17].

One drug that is particularly effective at targeting drug-induced adaptations in astrocytes is the antioxidant drug, N-acetylcysteine (NAC). NAC is a cystine pro-drug that has shown some promise clinically [18] but has been more extensively investigated preclinically as a research tool to reverse drug-mediated disruptions in astroglial glutamate homeostasis [15]. NAC has been shown to significantly limit heroin [19, 20], cocaine [21, 22], nicotine [23], and methamphetamine [14] seeking in rats with a drug self-administration history. Mechanistically, NAC is thought to prevent relapse by reversing drug-induced downregulation of GLT-1 in the NAcore. This hypothesis was first confirmed in studies focused on cocaine-mediated downregulated GLT-1 expression in the NAcore. In these studies, preventing NAC-mediated restoration of GLT-1 expression in the NAcore following cocaine SA and extinction blocked NAC’s ability to prevent relapse [10, 24]. While NAC does indeed facilitate extinction and prevent cued heroin seeking [19], the ability of NAC to normalize heroin-induced decreases in GLT-1 in the NAcore remains to be directly evaluated. However, additional drugs that upregulate GLT-1 expression (i.e. ceftriaxone) have been shown to normalize heroin-induced decreases in glutamate clearance in the NAcore, inhibiting relapse [11]. Importantly, ceftriaxone has also been shown to normalize cocaine-induced decreases in PAP association with synapses in the NAcore [13], suggesting a significant structure-function relationship underlying drug-induced decreases in PAP-synapse associations, alterations in glutamate clearance, and relapse.

While there is a substantial body of work regarding the consequences of heroin SA and extinction on neuronal and astroglial plasticity in the NAcore, little is known about the impact of heroin on neurons or astrocytes in the PrL cortex. This gap in our knowledge is particularly significant given that cue-induced glutamate release in the NAcore during heroin seeking requires activation of the PrL cortex [16]. While less is known about the relationship between PrL cortical astrocytes and opiates, there is some literature describing opiate-induced adaptations in PrL cortical astroglia. For example, chronic morphine treatment increases expression of glial fibrillary acidic protein (GFAP), a canonical marker for astrocytes [25], in the frontal cortex [26]. These data suggest that morphine induces astrogliosis, yet addition of morphine to astrocytes *in vitro* decreases proliferation and increases complexity of astrocyte processes, an effect prevented by naloxone pretreatment [27]. Thus, it is unclear how heroin SA and extinction may impact astrocyte structure in the PrL cortex and whether altered GFAP complexity is associated with altered astrocyte-mediated regulation of synaptic transmission.

Here we demonstrate that, in contrast to heroin-induced alterations in PAP interactions at NAcore synapses, heroin SA and extinction *increases* the association of PAPs with synaptic markers in the PrL cortex. This increased PAP-synaptic proximity was also accompanied by increased complexity of the GFAP cytoskeleton arbor within PrL cortical astrocytes. Importantly, both astroglial adaptations were normalized by repeated NAC treatment during extinction. Finally, heroin SA and extinction increased the size of dendritic spine heads yet decreased dendritic spine density in apical tuft dendrites of PrL-NAcore neurons. However, only the heroin-induced decreases in spine density were normalized by chronic NAC treatment. When investigating dendritic subcompartments on PrL-NAcore neurons, heroin-induced increases in dendritic spine head diameter were associated with increased GluA1 immunoreactivity in dendritic shafts and spine heads of PrL-NAcore neurons. Somewhat surprisingly, the increased GluA1 immunoreactivity in spine heads observed after heroin SA and extinction were further potentiated by NAC treatment. These results reveal that, in addition to normalizing astrocyte dysfunction in the NAcore, NAC normalizes heroin-induced astroglia and neuronal adaptations in the PrL cortex yet may lead to a potentiation of discrete inputs to the apical tuft of PrL-NAcore neurons.

## Materials and methods

### Animal subjects and surgery

Male Sprague-Dawley rats (*N*=63) were purchased from Charles Rivers Laboratory (Wilmington, MA) and were single-housed upon arrival in a temperature and humidity-controlled vivarium with standard rat chow (Harlan; Indianapolis, IN) and water available *ad libitum*. Rats were maintained on a 12-hr reverse light/dark cycle (lights off at 6 AM). All animal use protocols were approved by the Institutional Animal Care and Use Committee of the Medical University of South Carolina and were performed according to the National Institutes of Health Guide for the Care and Use of Laboratory Animals (8^th^ ed. 2011). At the time of surgery, all rats weighed 275-300g. On the day of surgery, rats were anesthetized with an intraperitoneal (i.p.) ketamine (66 mg/kg) and xylazine (1.33 mg/kg) injection, and received ketorolac (2.0 mg/kg, i.p.) for analgesia. A silastic catheter was implanted in the right jugular vein, attached to a cannula, which exited the animal’s back. Rats in Experiment 1 received a bilateral intra-PrL cortical (coordinates-AP: +2.8, ML: +/-0.6, DV: -3.8 mm from bregma) microinjection of AAV5-GfaABC1D-Lck-GFP (0.75 ul/hemisphere, Titer: 1×10^13^ vg/ml) to label the fine membrane processes of astrocytes [13, 28]. Rats in Experiment 2 received a bilateral microinjection of AAV1-CAG-Flex-Ruby2sm-FLAG (titer: 5×10^12^ vg/ml) as well as intra-NAcore (coordinates-AP: +1.7, ML: +/- 1.6, DV: -7 mm relative to bregma) injection of rgAAV-hSyn-Cre-WPRE (0.75 ul/hem, titer: 7×10^12^) to specifically target PrL cortical-NAcore neurons for dendritic spine analyses [29]. Injections (0.75 ul/hemisphere) were performed over a period of 5 minutes (0.15 ul/minute) using a Nanoject II (Drummond Scientific, Broomall, PA) and injectors were left in place for 5 minutes to facilitate viral diffusion from the injection site. All constructs were purchased from Addgene (Cambridge, MA) except AAV5-GfaABC1D-Lck-GFP which was custom-packaged at the University of North Carolina Gene Therapy Center viral vector core (Chapel Hill, NC). Rats were allowed at least 5 days of post-operative recovery during which catheters were flushed daily with taurolidine citrate solution (TCS, Access Technologies, Skokie, IL) and food and water were available *ad libitum*. For each experiment, ≥2 independent cohorts were used.

### Drugs

Heroin hydrochloride was provided by the National Institute on Drug Abuse. N-acetylcysteine was purchased from Sigma (#A7250) and diluted in sterile saline to 100 mg/ml.

### Heroin self-administration and extinction procedures

All experiments were performed between the hours of 8AM and 6PM. All SA experiments were performed in standard MedPC operant chambers equipped with two retractable levers, a tone generator, and a cue light (Med Associates Fairfax, VT). The beginning of each session was signaled by illumination of the houselight and lever availability. Rats were trained for 14 days (3 hrs/day) to lever press for heroin on a fixed ratio 1 (FR1) schedule of reinforcement. On days 1-2, active lever presses (ALP) resulted in a light and tone cue delivery followed by a 100 μg/50 μl iv bolus of heroin hydrochloride dissolved in sterile saline. On days 3-4, rats received a 50 μg/50 μl bolus followed by a 20 sec. timeout period, and on days 5-14 rats received a 25 μg/50 μl bolus (maintenance dose). Yoked-saline animals were paired to a heroin SA animal; receiving non-contingent saline infusions in a temporally-paired manner. SA criteria were set at ≥1 infusion per session on days 1-4 whereas animals were required to receive ≥10 infusions per session on days 5-14. Following SA, all rats entered extinction whereby ALP resulted in no cue or drug delivery. All rats received 12-13 days of extinction training (3 hours/day). Beginning on day 4, a subset of animals received N-acetylcysteine (100 mg/kg, i.p.) or vehicle (0.9% NaCl) 30 minutes before each subsequent session, and all animals injected with N-acetylcysteine received exactly 13 days of extinction training. All yoked-saline animals received vehicle injections during extinction. This dose of NAC has previously been shown to reduce extinction responding on the active lever and provide an enduring reduction in cue- and primed-heroin seeking using the same heroin SA protocol [19].

### Transcardial perfusions and immunohistochemistry

Twenty-four hours after the final extinction session, rats were transcardially perfused with 150 ml of 0.1M phosphate buffer (PB), followed by 200 ml of 4% paraformaldehyde (PFA) in 0.1M PB. Brains were extracted and post-fixed in PFA for 24 hours. Sections (80 µms thick) containing the PrL cortex and NAcore were collected in 0.1M PBS containing 0.02% sodium azide with a vibrating microtome (Leica), and stored at 4ºC until processing.

Immunohistochemistry (IHC) was performed as previously described [13, 14, 30]. Briefly, 3 sections per animal were blocked with 0.1M PBS containing 2% (v/v) TritonX-100 (PBST) containing 2% (v/v) normal goat serum (NGS). Sections were then incubated overnight with gentle agitation at 4ºC with the following primary antisera: Experiment 1 – chicken anti-GFP (Abcam, 1:1000, ab13970 RRID:AB_300798) + mouse anti-GluA2 (Millipore, 2.13 ug/ml, MAB397 RRID:AB_11212990) or rabbit anti-Synapsin I (Abcam, 1:500, ab64581 RRID:AB_1281135), and in a separate run, rabbit anti-GFAP (Abcam, 1:1000, ab7260 RRID:AB_305808); Experiment 2 – mouse anti-FLAG (Sigma, 1:2000, F1804 RRID:AB_262044) and rabbit anti-GluA1 (Abcam, 1:500, ab31232 RRID:AB_2113447). Sections were then washed 3×5 minutes with PBST, and were incubated in species-specific secondary antisera (Invitrogen, all 1:1000) conjugated to Alexa Fluor 488 (RRID:AB_2534096, GFP), 594 (RRID:AB_2534091, FLAG), or 647 (RRID:AB_2535804, GFAP, Synapsin I, GluA1 and GluA2) diluted in PBST with 2% NGS for 5 hours at room temperature with gentle agitation. Sections were washed 3×5 minutes in PBST, then mounted on Superfrost plus slides with Prolong Gold antifade (ThermoFisher Scientific). Slides were stored at 4ºC until imaging (< 2 months).

### Confocal microscopy

For all high-resolution confocal microscopy experiments, a Leica SP8 confocal microscope equipped with HyD detectors for enhanced sensitivity was used. Laser power, gain, and pinhole size (≤1.5 Airy Unit) were empirically determined to obtain optimal saturation values and subsequent deconvolution results on a per-experiment basis. Laser power and gain values were then held relatively constant for the remainder of each experiment. For all experiments, Alexa488-mediated Lck-GFP detection was carried out using an Argon 488nm laser, Alexa-594-mediated FLAG detection was carried out using an OPSL 552nm laser, and Alexa647-mediated GluA2, GFAP, and Synapsin I detection was carried out using a Diode 638nm laser. In experiment 1a, fields of GFAP+ cells were imaged in layers II/III and V of the PrL cortex at a 4096×4096 frame size, 0.1 μm Z-step size, 1X digital zoom, with a line average of 4 using a 63X oil-immersion objective (1.4 N.A.). In experiment 1b, individual Lck-GFP-expressing astrocytes that were visually isolated from neighboring astrocytes in layers II/III and V as well as GluA2/Synapsin I puncta were sampled using a 63X oil-immersion objective (1.4 N.A.) at a 2048×2048 frame size, 0.3 μm Z-step size, 1.5X digital zoom, with a frame average of 4. Care was taken to ensure that the majority of each astrocyte in the Z-plane was imaged. In experiment 2, apical tuft dendrites of layer V PrL-NAcore neurons were imaged as previously described [30]. Briefly, dendritic spine segments were imaged only if they met the following criteria: 1: they could be traced back to the cell body of origin, 2: they were unobscured by neighboring dendritic segments, and 3: they were past the bifurcation of the proximal dendrite (terminating in layers I and II). Dendritic spine segments (∼50 μm) were collected at 1024×512 frame size, 0.1 μm Z-step size, 4.1X digital zoom, with a line average of 2. Each spine segment imaged contained a GluA1 channel in addition to FLAG for subsequent co-registry analyses.

### Image analysis

All image analyses were performed by an investigator blind to experimental groups. Following acquisition, confocal Z-stacks were deconvolved using Huygens software (Scientific Volume Imaging, Hilversum, NL) and then exported to Bitplane Imaris (v. 9.0.1) for digital rendering and analysis. Complexity of GFAP arbors were analyzed as previously described[14]. Briefly, Z-stacks were cropped to 12 μm in the Z plane to identify fields of cells with homogenous signal intensity, avoiding large areas of neurovasculature. We used a previously published semi-automated approach to skeletonize GFAP arbors belonging to individual, non-overlapping astrocytes. In doing so, we set the minimum diameter of a GFAP filaments to 1 μm. Iterative processing of GFAP skeletons is then performed in order to accurately skeletonize each astrocyte within the field. Once completed, we exported and analyzed astrocyte structural variables using the following outputs from the Imaris filament function: the number of astrocytes in each image, the total sum of the GFAP filament length for each cell per image, the average GFAP diameter of each cell per image, the average number of branch points for each cell per image, and the average number of 3D Sholl’s intersections (1 µm radius) per cell, per image. The average number of Sholl’s intersections was then binned by 5 µm radii from 0-50. Data is expressed as an average for each cell, or an overall Z-series data set average as specified in the results.

The whole-cell astrocyte synaptic contact assays were performed as described previously [13, 14], with minor modifications. Briefly, confocal Z-stacks were cropped in 3D to isolate each astrocyte. A 3D space-filling model was generated from the Lck-GFP signal from each virally labeled astrocyte. We then masked out all signal outside of each astrocyte region of interest (ROI) to eliminate background. The colocalization module of Imaris was used to perform co-registry analyses. A signal intensity threshold was automatically set according to the Costes method [31] for each channel (Channel 1: GFP, Channel 2: Synapsin I or GluA2). Images that could not be assigned automatic thresholds, due to heterogenous signal intensity distribution through the Z-plane of the dataset, were not analyzed. Voxels that contained signal above threshold for each channel and occupied the same physical space were considered co-registered. Within this assay a greater number of co-registered voxels indicates a greater degree of astrocyte association at synapses on a whole-cell level. A 3D space-filling model was then built on the channel containing co-registered voxels. The total volume of this channel was then normalized to the total volume of the isolated astrocyte to generate a %ROI co-registered measurement. We also digitally modeled each Synapsin I or GluA2 puncta to determine the effect of treatment on the number of puncta within each image, normalized to the dataset volume. Data exported included the %ROI co-registered normalized to the saline control group (% co-registered), the ratio of the astrocyte surface area to volume, and the number of Synapsin I/GluA2 puncta normalized to the dataset volume.

Dendritic spine morphometric feature and GluA1 co-registry were analyzed as previously described [30]. Briefly, the filament extension of Imaris was used to manually trace each dendritic shaft and semi-automatically label corresponding dendritic spines. Care was taken to ensure that each dendritic spine head was modeled accurately. Variables exported included the average dendritic spine head diameter (d_H_) of each segment and the dendritic spine density (spines/µm). We also binned the spine head diameters (0.2 µm bins) to generate a frequency distribution. For GluA1 co-registry analyses, a 3D space-filling model was then used to isolate the dendrite segment (shaft + spines), and GluA1 signal above background contained within the dendrite was isolated. The colocalization module of Imaris was then used to generate a channel containing GluA1 and FLAG co-registry and the ‘filament analysis’ MATLAB Xtension was used to split the channel into FLAG-GluA1 co-registry in dendritic subcompartments including the dendritic spine head and dendritic shaft. Output variables included the percent of GluA1 volume normalized to the dendrite volume, the total volume of spine head-specific GluA1 normalized to the dendrite volume, and the total volume of dendritic shaft GluA1 normalized to the dendrite volume as well as signal intensity controls. Data is expressed as animal averages; collapsing all dendritic spine morphometric data across segments sampled from each animal.

### Experimental design

**Figure 1A** shows the experimental timelines. Experiment 1 was designed to examine the effects of heroin self-administration and extinction, with or without NAC administration, on GFAP complexity in the PrL cortex (1a) as well as astrocyte-synapse proximity (1b). Experiment 2 was designed to examine whether dendritic spine morphometric properties, and associated GluA1 immunoreactivity in discrete dendritic sub compartments, were altered in PrL-NAcore neurons after extinction of heroin SA with or without NAC administration.

**Figure 1.**
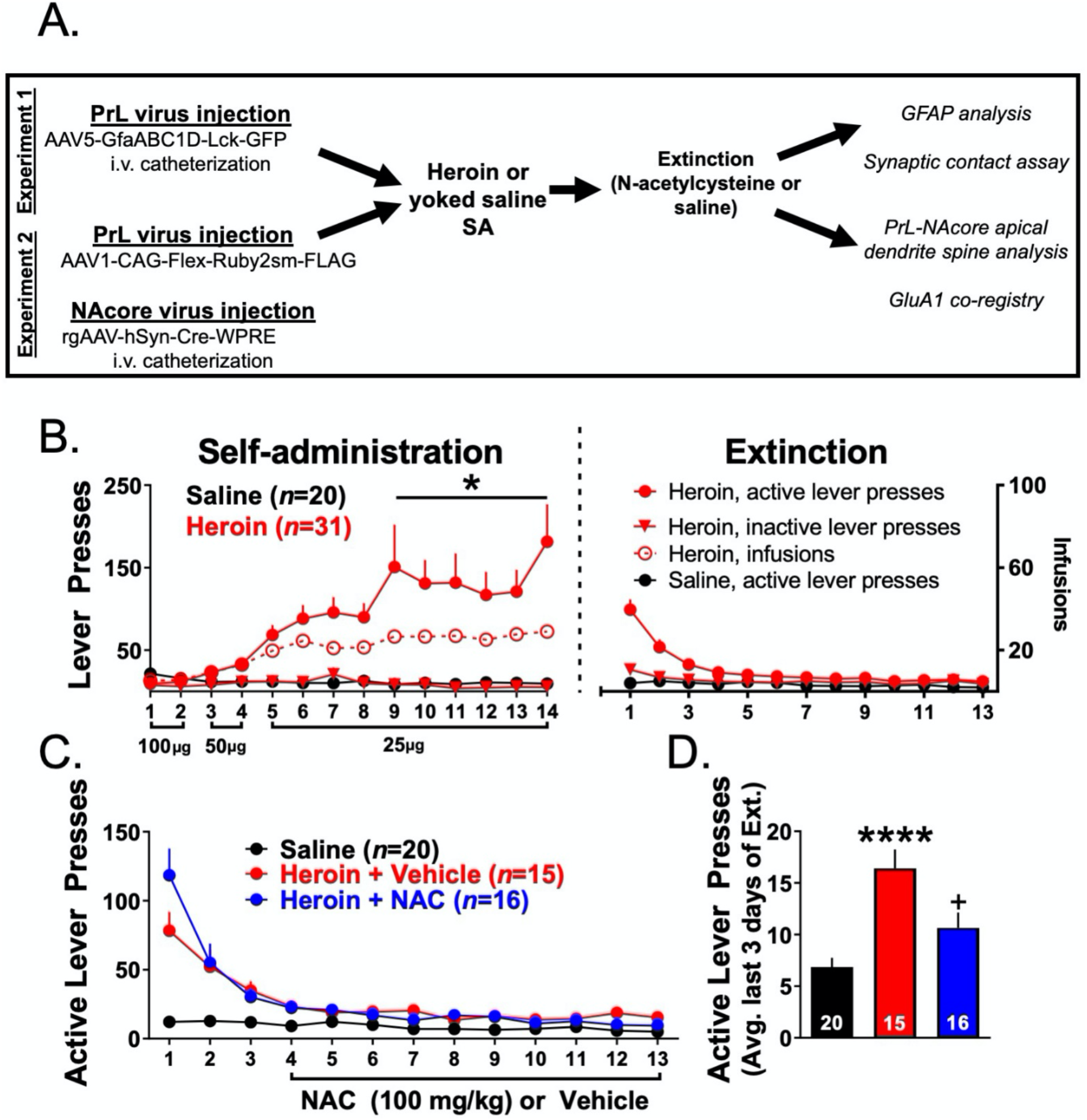
NAC reduces extinction responding. A. Diagram illustrating the experimental timeline for each experiment. B. Self-administration and extinction behavior for heroin and Yoked-Saline rats. Lever presses are shown on the left y axis and infusions are shown on the right y axis. \**p*<0.05 compared to saline active lever presses. C. Active lever presses during extinction for Yoked-Saline, heroin animals injected with vehicle, and heroin animals injected with NAC. D. Average active lever presses over the last 3 days of extinction for the three experimental groups. \*\*\*\**p*<0.0001 compared to Yoked-Saline, +*p*<0.05 compared to Heroin + Vehicle. The number of animals within each group are shown within the bars.

### Statistical analyses

All statistical analyses were performed with Graphpad Prism software (v. 8.0, La Jolla, CA). When analyzing three groups across a single dependent variable, a one-way ANOVA was performed followed by Tukey’s multiple comparison test if a significant effect of treatment was revealed. Normality of data within each group was analyzed using Shapiro-Wilk test. If one or more groups for each variable analyzed did not pass the Shapiro-Wilk test, a non-parametric Kruskal-Wallis test was used. In this instance, post hoc tests controlled for the false discovery rate (FDR) using the two-stage linear step-up procedure of Benjamini, Krieger,and Yekutieli if a significant effect was revealed. If data were sampled from a normal distribution, but between group standard deviations were not equal, a Brown-Forsythe-corrected one-way ANOVA was performed followed by Dunnett’s T3 test when a significant effect of treatment was revealed. When three groups were compared across a repeated measure, a repeated measure one-way ANOVA was applied followed by Tukey’s multiple comparison test comparing the mean of the repeated measure between groups. Dendritic spine frequency distribution data as well as self-administration data were analyzed with a two-way repeated measure ANOVA followed by Tukey’s multiple comparison test when a significant interaction was revealed. Unless otherwise stated, each data point represents an animal average derived from multiple images or SA/extinction sessions. Statistical outliers were detected according to the ROUT method (Q=1%) and these data were removed from analyses as specified in the results. All data is expressed as the mean +/- SEM and significance was set at *p*<0.05.

## Results

### Heroin self-administration and extinction: Experiments 1 and 2

Of the 63 initial animals in Experiments 1 and 2, 12 animals did not finish SA due to catheter failure. **Figure 1B** shows self-administration and extinction data for heroin SA (*N*=31) and Yoked-Saline (*N*=20) rats used for all experiments. When comparing heroin and Yoked-Saline active lever presses across SA sessions, a two-way repeated measures ANOVA revealed a significant group by time interaction (F(13,367)=5.11, *p*<0.0001). Bonferroni’s multiple comparison test revealed that heroin SA animals pressed the active lever significantly more than Yoked-Saline controls in sessions 9-14 (*p*<0.05). There was no difference between groups (that were subsequently divided by treatment during extinction) in the average number of heroin infusions earned over the last 3 days of SA (Heroin + Vehicle: 29.38 +/-2.80, Heroin + NAC: 25.75+/-4.87). **Figure 1C** shows extinction training data for Yoked-Saline animals injected with Vehicle (*n*=20), and heroin SA rats injected with Vehicle (*n*=15) or NAC (*n*=16) during the last 10 days of extinction. **Figure 1D** shows active lever presses averaged across the last 3 days of extinction for each group. A one-way ANOVA revealed a significant effect of treatment (F(2,48)=12.15, *p*<0.0001). Tukey’s multiple comparison tests revealed that heroin SA animals showed significantly more lever pressing during the last 3 days of extinction than Yoked-Saline rats (*p*<0.0001) and that NAC significantly decreased lever pressing to Yoked-Saline values (*p*=0.019).

### Experiment 1A – Heroin SA and extinction increased astroglial GFAP arbor complexity, an effect that was prevented by repeated NAC treatment during extinction

Of the initial 10 Yoked-Saline rats used in this experiment, 2 were not included due to technical issues related to IHC processing, 2 Heroin + Vehicle rats were not included due to technical issues related to IHC, and one Heroin + NAC animal was excluded as a result of inadequate immunohistochemical labeling of GFAP. **Figure 2A** shows a representative field of GFAP+ cells in the PL cortex (left); the same image is shown with an overlaid Imaris-based skeletonization (white) of each GFAP+ cell (middle), and an individual GFAP+ cell (yellow) isolated from the other cells (right). Using this high throughput approach, we analyzed GFAP complexity in 3-7 Z-series data sets from 8 Yoked-Saline rats, 7 Heroin + Vehicle rats, and 9 Heroin + NAC rats using several different metrics. We first analyzed the average number of GFAP+ cells per field. A one-way ANOVA revealed no effect of treatment (F(2,92)=1.23, *p*=0.30, **Figure 2B)**. For complexity analyses, we first analyzed the total sum of GFAP filament length within each Z-series data set **(Figure 2C)**. A Shapiro-Wilk test indicated that data in Yoked-Saline (W=0.9, *p*=0.0005) and Heroin + NAC (W=0.94, *p*=0.04) groups were non-normally distributed. Accordingly, a Kruskal-Wallis non-parametric ANOVA was used which indicated a significant difference between groups (K-W=12.99, *p*=0.002). The two-stage linear step-up procedure revealed that heroin increased total GFAP filament length per image when compared to Yoked-Saline (*q*=0.0007), which was normalized by NAC treatment during extinction (*q*=0.004). We next analyzed the average GFAP filament diameter for each cell per z-series data set **(Figure 2D)**. One Z-series data set from Yoked-Saline and 2 Z-series data sets from the Heroin + NAC group were identified as statistical outliers. A Brown-Forsythe test revealed that the standard deviations were significantly different between groups (F(2,89)=3.26, *p*=0.04). As such, these data were analyzed using a Brown-Forsythe-corrected ANOVA which revealed there was no significant effect of treatment (F(2,74.38)=0.626, *p*=0.54). We next analyzed the average number of branch points of each cell, within each z-series data set **(Figure 2E)**. One Z-series data set from the Yoked-Saline group was identified as a statistical outlier. A Shapiro-Wilk test indicated that data from the Yoked-Saline group was non-normally distributed. Accordingly, a Kruskal-Wallis test was used which revealed a significant difference between groups (K-W=14.61, *p*=0.0007). The two-stage linear step-up procedure revealed that heroin increased the average number of branch points within GFAP filament arbors compared to Yoked-Saline (*q*=0.0002) which was normalized by NAC treatment during extinction (*q*=0.003). We also performed a 3D Sholl’s analysis for each cell within each image using 1 µm 3D concentric sphere radii (which were then compiled into 5 µm bins) and analyzed the average number of intersections as a function of distance from the center of each cell (**Figure 2F**). A repeated measures one-way ANOVA revealed a significant difference between group means (F(2,20)=19.23, *p*<0.0001). Tukey’s multiple comparison test revealed that heroin increased the average number of Sholl’s intersections compared to Yoked-Saline (*p*=0.0002), which was normalized by NAC treatment during extinction (*p*<0.0001). In a subsequent analysis, we focused on the first 15 µm from the center of each cell (**Figure 2G**). In these data, a one-way ANOVA revealed a significant difference between groups (F(2,92)=8.41, *p*=0.0004). Tukey’s multiple comparison test revealed that heroin increased the number of Sholl’s intersections in this region when compared to Yoked-Saline (*p*=0.003), which was normalized by NAC treatment during extinction (*p*=0.0007). **Figure 2H** shows an individual cell with a representative skeletonized GFAP arbor from each treatment group.

**Figure 2.**
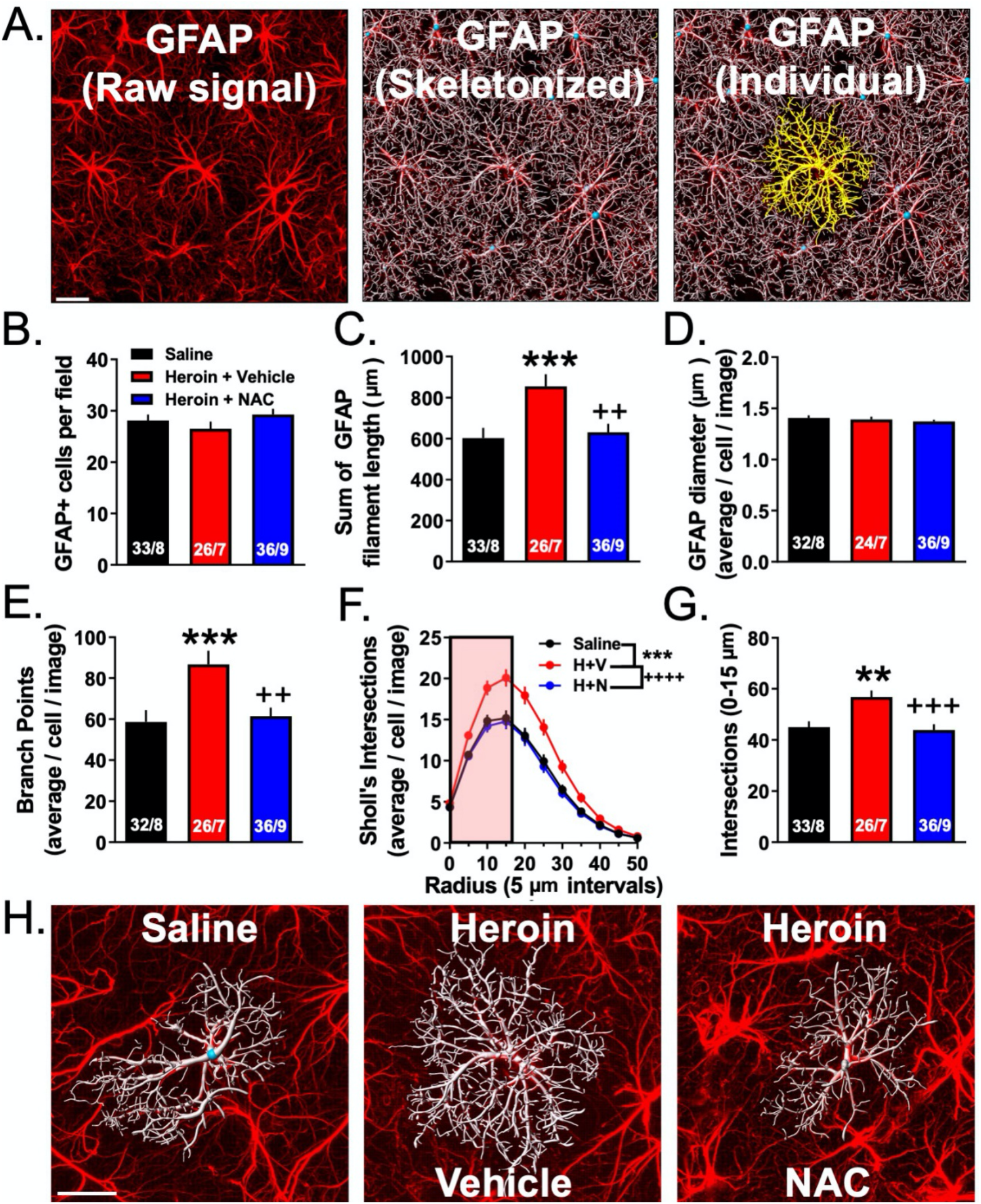
NAC normalizes heroin-induced alterations in PrL cortical astrocyte complexity. A. Representative field of GFAP+ astrocytes skeletonized for analysis. Left-Raw GFAP signal from the confocal microscope. Middle–Skeletonized GFAP. Right-Isolated GFAP+ astrocyte from the field of cells. B. There was no difference between groups in the number of GFAP+ cells per field. C. Heroin increased the total length of the GFAP cytoskeletal filament which was normalized by repeated NAC treatment. D. There was no difference between groups in the average diameter of the GFAP filament. E. Heroin increased the average number of branch points which was normalized by repeated NAC treatment. F. Frequency distribution of the number of Sholl’s intersections as a function of distance from the center of the cell. Heroin increased the number of Sholl’s intersections averaged across all bins which was normalized by repeated NAC treatment. G. Average number of Sholl’s intersections across the first 4 bins (0, 5, 10, and 15 µm). Heroin increased the number of Sholl’s intersections which was normalized by chronic NAC treatment. H. Representative skeletonized GFAP+ cells for each treatment group. \*\**p*<0.01, \*\*\**p*<0.001 compared to Yoked-Saline, ++*p*<0.01, +++*p*<0.001, ++++*p*<0.0001 compared to Heroin + Vehicle. Numbers within each bar denote the number of images taken / the number of animals used. Scale bars indicate 20 µm.

### Experiment 1B – Heroin SA and extinction increased astrocyte association with multiple synaptic markers; an effect that was prevented by repeated NAC treatment during extinction

Here we used an astrocyte-specific membrane-targeted GFP reporter virus to label the astroglial plasma membrane (AAV5-GfaABC1D-Lck-GFP, **Figure S1)**. Using this construct, we performed two separate astrocyte-synaptic contact analyses using PrL cortical sections collected from the three experimental groups: Yoked-Saline (*n*=10), Heroin + Vehicle (*n*=9), and Heroin + NAC (*n*=10). In the first set of contact assays, astroglial structure was visualized with viral vector-mediated membrane targeted GFP expression achieved with the AAV5-GfaABC1D-Lck-GFP (Lck-GFP) viral vector and further amplified with GFP IHC and excitatory synapses were labeled with GluA2 IHC as described previously [32]. The second set of astrocyte-synaptic contact analyses was performed identically, except that Synapsin I IHC was used as a general synaptic marker [13]. We imaged 3-7 cells/animal for GluA2 and 3-5 cells/animal for Synapsin I. For the GluA2 analyses, the following were excluded: 1 Yoked-Saline animal was missing localization of Lck-GFP virus expression, 1 Yoked-Saline animal for issues related to IHC, 1 yoked saline animal for lack of cells satisfying the Costes threshold assignment, 1 Heroin + Vehicle animal had insufficient Lck-GFP virus expression, 1 Heroin + NAC animal for lack of cells satisfying Costes threshold detection, and 1 Heroin + NAC animal had no more available tissue.

**Figure 3A** shows a representative PrL cortical Lck-GFP-labeled astroglia (green) and punctate signal corresponding to GluA2 (red). To aid in the visualization of the punctate GluA2 synaptic marker signal and the convoluted astroglial plasma membrane, an individual optical section within the center of an astrocyte was selected from the many images that comprise each Z-series data to provide visual detail. The left panel shows the raw signal, the middle panel shows GluA2 signal contained within the 3D surface of the astrocyte membrane, and the right panel depicts co-registry of the two signals. Within each panel, a smaller depiction of the full z-series data set is visible in the top right corner, with a white box depicting the location of the individual optical section seen within the single section view. **Figure 3B** shows the percent of the co-registry of the GFP and GluA2 across the three treatment groups, this output is used as a metric of astrocyte-synaptic interaction by our lab and others, for more information see [13, 14, 33, 34]. To facilitate comparison of our experimental measurements utilizing both a general synaptic marker and an excitatory synaptic marker, all data were normalized to Yoked-Saline and expressed as % of control. This facilitates comparison between markers and cohorts of animals. However, it is important to note that despite the fact that baseline values differed slightly from cohort to cohort, the effect of heroin was indeed reproducible across both cohorts (**Figure S2A-D**). For GluA2, a Brown-Forsythe-corrected ANOVA revealed a significant effect of treatment (F(2,9.743)=4.74, *p*<0.01). The two-stage linear step-up procedure indicated that the Heroin + Vehicle group showed increased co-registry of GluA2 and GFP compared to the Yoked-Saline group (*p*=0.01), which was normalized by repeated NAC treatment during extinction (*p*<0.02). We next determined whether this effect was due to an increased number of GluA2 puncta within each dataset. A one-way ANOVA revealed no significant effect of treatment (F(2,20)=2.24, *p*=0.13, **Figure 3C). Figure 3D** shows representative 3D-reconstructed astroglia for each treatment group with GluA2 puncta (top) as well as the extent of synaptic contact as visualized by white GFP and GluA2 co-registered voxels (bottom).

**Figure 3.**
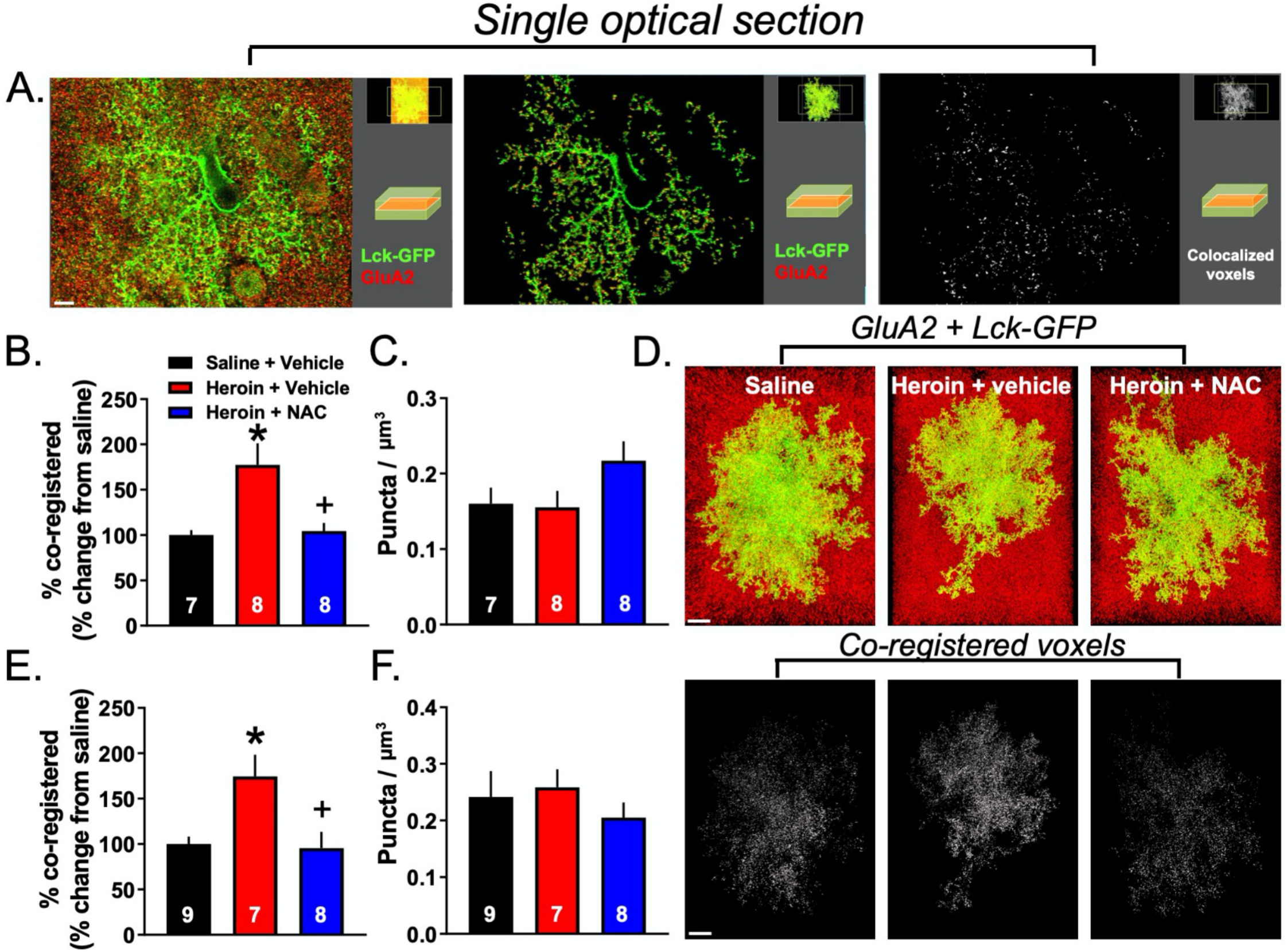
NAC normalized heroin-induced increases in PrL cortical astrocyte PAPs-synapse association. A. A single optical section from a representative Lck-GFP-labeled astrocyte and GluA2 puncta. Left-Raw signal from confocal microscope. Middle-GluA2 confined within the proximity of the astrocyte membrane. Right-co-registered voxels within the proximity of the astrocyte membrane. B. Heroin SA and extinction increased the percent co-registration of GFP and GluA2 which was normalized by repeated NAC treatment. C. There was no difference between groups in the number of GluA2 puncta normalized to the dataset volume. D. Representative astrocytes from the three groups. Top-Raw signal from the confocal microscope. Bottom-co-registered voxels within the region occupied by each astrocyte. E. Heroin SA and extinction increased the percent co-registration of GFP and Synapsin I which was normalized by repeated NAC treatment. F. There was no difference between groups in the number of Synapsin I puncta normalized to the dataset volume. Numbers within each bar denote the number of animals used. \**p*<0.05 compared to Yoked-Saline, +*p*<0.05 compared to Heroin + Vehicle. Scale bars indicate 5 µm (A) and 10 µm (D).

As described above, we also performed the same imaging and analysis for Lck-GFP and Synapsin I, using tissue from the same animals to determine whether the observed effect was specific to synapses containing the GluA2 subunit of AMPA receptors. 1 Yoked-Saline animal, two Heroin + Vehicle animals, and two Heroin + NAC animals were removed from the analysis for the following reasons: Yoked-Saline + Vehicle: off-target Lck-GFP virus expression (*n*=1); Heroin + Vehicle: lack of available tissue (*n*=1) or insufficient Lck-GFP virus expression (*n*=1); Heroin + NAC: lack of a sufficient number of analyzable cells after imaging (*n*=1), and technical issues related to IHC (*n*=1). A one-way ANOVA revealed a significant main effect of treatment (F(2,21)=6.51, *p*=0.006, **Figure 3E**). Tukey’s multiple comparison test revealed that heroin SA and extinction increased astrocyte-synaptic interaction as assayed by Lck-GFP-Synapsin 1 co-registration when compared to Yoked-Saline controls (*p*=0.012), which was normalized by NAC treatment during extinction (*p*=0.012). Akin to analyses performed for GluA2, we also analyzed the number of Synapsin I voxels in a subset of images from each animal **(Figure 3F)**. Shapiro-Wilk’s test indicated that data from Yoked-Saline controls (W=0.83, *p*=0.04) were non-normally distributed. A Kruskal-Wallis test indicated no difference between groups (K-W=2.04, *p*=0.36). There was no difference between groups in astrocyte complexity as measured by the ratio of the surface area to volume for cells used in either GluA2 or Synapsin I data sets **(Figure S2 E-F)**.

### Experiment 2 – Heroin SA and extinction impacts spine density and spine head diameter on PrL-NAcore neurons; an effect that was partially reversed by repeated NAC treatment during extinction

Others have shown that heroin SA decreases the number of dendritic spines on PrL cortical neurons after one month of abstinence [35]. Here, we sought to determine whether heroin SA and extinction altered dendritic spine morphometrics specifically in PrL cortical neurons projecting to the NAcore. We imaged 4-8 dendritic spine segments from Yoked-Saline (*n*=10), Heroin + Vehicle (*n*=6) and Heroin + NAC (*n*=6) animals after extinction. **Figure 4A** shows a representative layer V PrL-NAcore neuron fully labeled with the highlighted area indicating the region sampled at the apical tuft. **Figure 4B** shows the dendritic spine segment sampled from the highlighted region in 5A. **Figure 4C** shows the filament-processed segment for spine detection and **Figure 4D** shows a color-coded map of dendritic spine d_H_, with warmer colors indicating larger values. We first analyzed the average dendritic spine d_H_ across the three treatment groups. A one-way ANOVA revealed a significant effect of treatment (F(2,19)=20.70, *p*<0.0001, **Figure 4E**). Tukey’s multiple comparison test revealed that heroin SA and extinction increased the average dendritic spine d_H_ (*p*<0.0001), which was not altered by NAC treatment during extinction (*p*=0.0007). Next, we investigated whether this effect was specific to smaller or larger spines. To do so, we binned the spine d_H_ (0.2 µm bins) and generated a frequency distribution. A two-way repeated measures ANOVA revealed a significant treatment by bin interaction (F(8,76)=8.144, *p*<0.0001, **Figure 4F**). Tukey’s multiple comparison test revealed that heroin SA and extinction, regardless of NAC treatment, decreased the proportion of smaller spines (<0.2 – *p*<0.0001; 0.2-0.4 – *p*<0.05), but increased the proportion of spines with larger d_H_ (0.4-0.6 – *p*<0.01) compared to Yoked-Saline rats. Interestingly, the proportion of spines with d_H_ between 0.6-0.8 was increased in the Heroin SA + Vehicle group (*p*=0.008) but there was no effect of NAC on this specific bin compared to Yoked-Saline (*p*=0.17). We next analyzed the average dendritic spine density in all three treatment groups from the same dendrite segments. A one-way ANOVA revealed a significant effect of treatment (F(2,19)=13.46, *p*=0.0002, **Figure 4G**). Tukey’s multiple comparison test indicated that heroin SA and extinction decreased the average spine density compared to saline controls (*p*=0.0003), which was prevented by NAC treatment during extinction (*p*=0.002). To determine which spine head diameter bins accounted for the altered spine density, we performed a similar analysis as in Figure 5F, but instead normalized the number of spines within each bin to length of the dendrite. A two-way repeated measures ANOVA revealed a significant treatment by bin interaction (F(8.76)=8.722, *p*<0.0001, **Figure 4H**). Tukey’s multiple comparison test indicated that heroin SA and extinction (*p*<0.0001), regardless of NAC treatment (*p*=0.0003), decreased the density of spines with a head diameter less than 0.2 µm. However, the Heroin + Vehicle group showed decreased spines in the 0.2-0.4 µm bin compared to Yoked-Saline (*p*<0.0001), which was normalized in the Heroin + NAC group (*p*<0.0001 compared to Heroin + Vehicle). Interestingly, the density of larger spines with head diameters between 0.4-0.6 µm were unaffected in the Heroin + Vehicle group compared to Yoked-Saline (*p*=0.194), yet Heroin + NAC increased the density of these spines compared to Yoked-Saline (*p*=0.0125) and Heroin + Vehicle (*p*=0.0002) groups. These effects on dendritic spine morphometric properties were likely not due to inherent differences in signal intensity within the spine heads as there was no difference between groups in the average FLAG intensity within the spine head region **(Figure S3A)**. Representative spine segments (15 µm cropped segments) are shown in **Figure 4I**.

**Figure 4.**
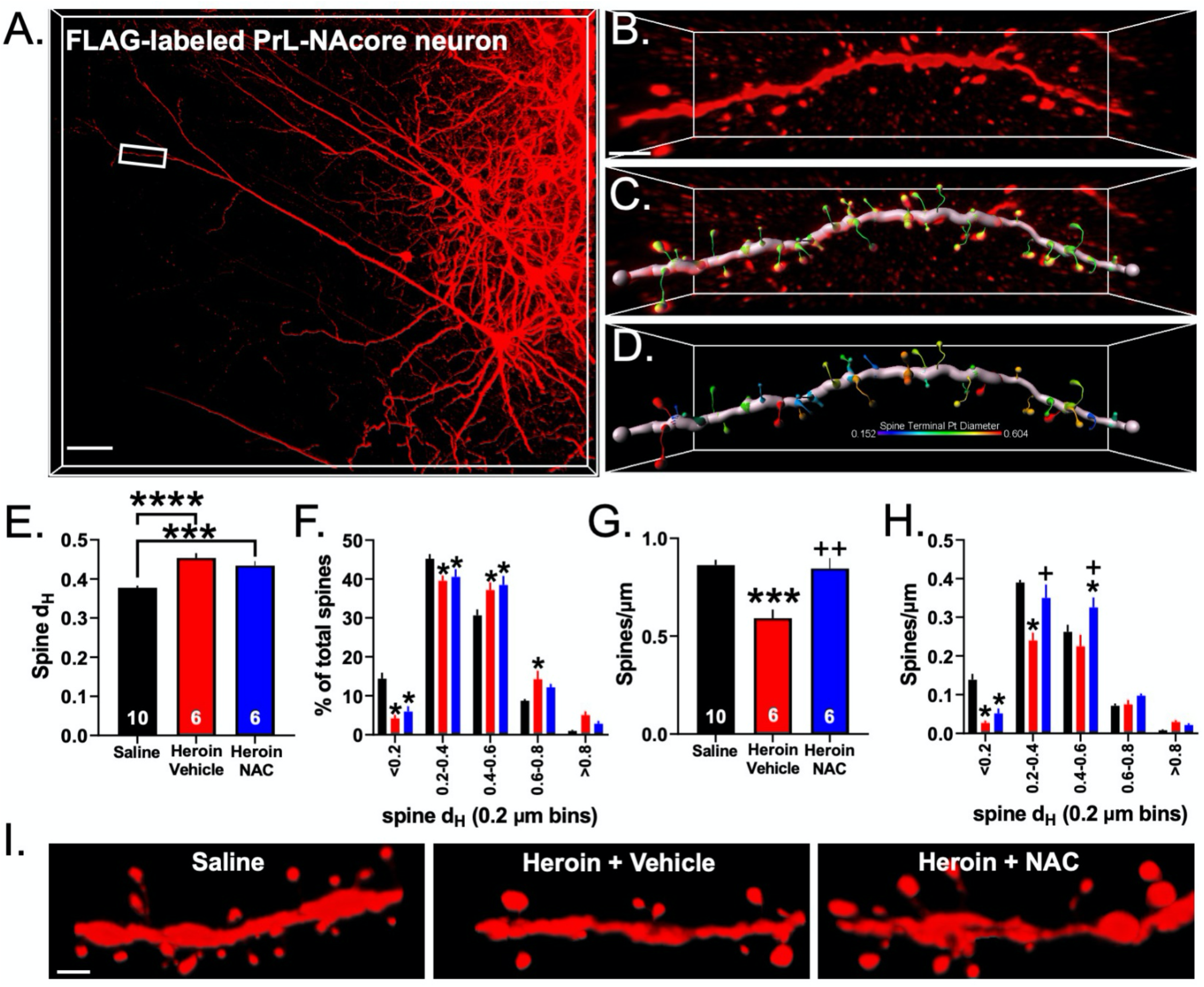
NAC normalized heroin-induced decreases in PrL-NAcore apical spine density, but not head diameter. A. Left-Representative FLAG-labeled PrL-NAcore neuron. B-D. B) Dendritic spine segment taken from the apical tuft (boxed region in A), C) Filament processed spine segment, D) Heat map of dendritic spine head diameters along the segment – warmer colors indicate a greater head diameter. Note that most spines fall within the middle range of head diameters. E. Heroin SA and extinction increased the average spine head diameter of PrL-NAcore neurons which was not affected by repeated NAC treatment. F. Frequency distribution of the percent of total spines as a function of spine head diameter bin (0.2 µm). Heroin SA and extinction regardless of NAC treatment decreased the proportion of spines with smaller head diameters and increased the proportion of spines with larger head diameters. G. Heroin SA and extinction reduced the number of spines per µm of dendrite which was normalized by repeated NAC treatment during extinction. H. Heroin SA and extinction regardless of NAC treatment reduced the density of spines with smaller head diameters, yet NAC treatment normalized a heroin-induced decrease in spines with a head diameter between 0.2-0.4 µm. NAC treatment during extinction from heroin SA also increased the density of spines with a head diameter between 0.4-0.6 µm. I. Representative spines from a 15 µm stretch of dendrite are shown. d_H_=head diameter. Numbers within each bar denote the number of animals used. \**p*<0.05, \*\**p*<0.01, \*\*\**p*<0.001, \*\*\*\**p*<0.0001 compared to Yoked-Saline. +*p*<0.05, ++*p*<0.01 compared to Heroin + Vehicle. Scale bars indicate 70 µm (A), 3 µm (B-D), and 2 µm (I).

**Figure 5.**
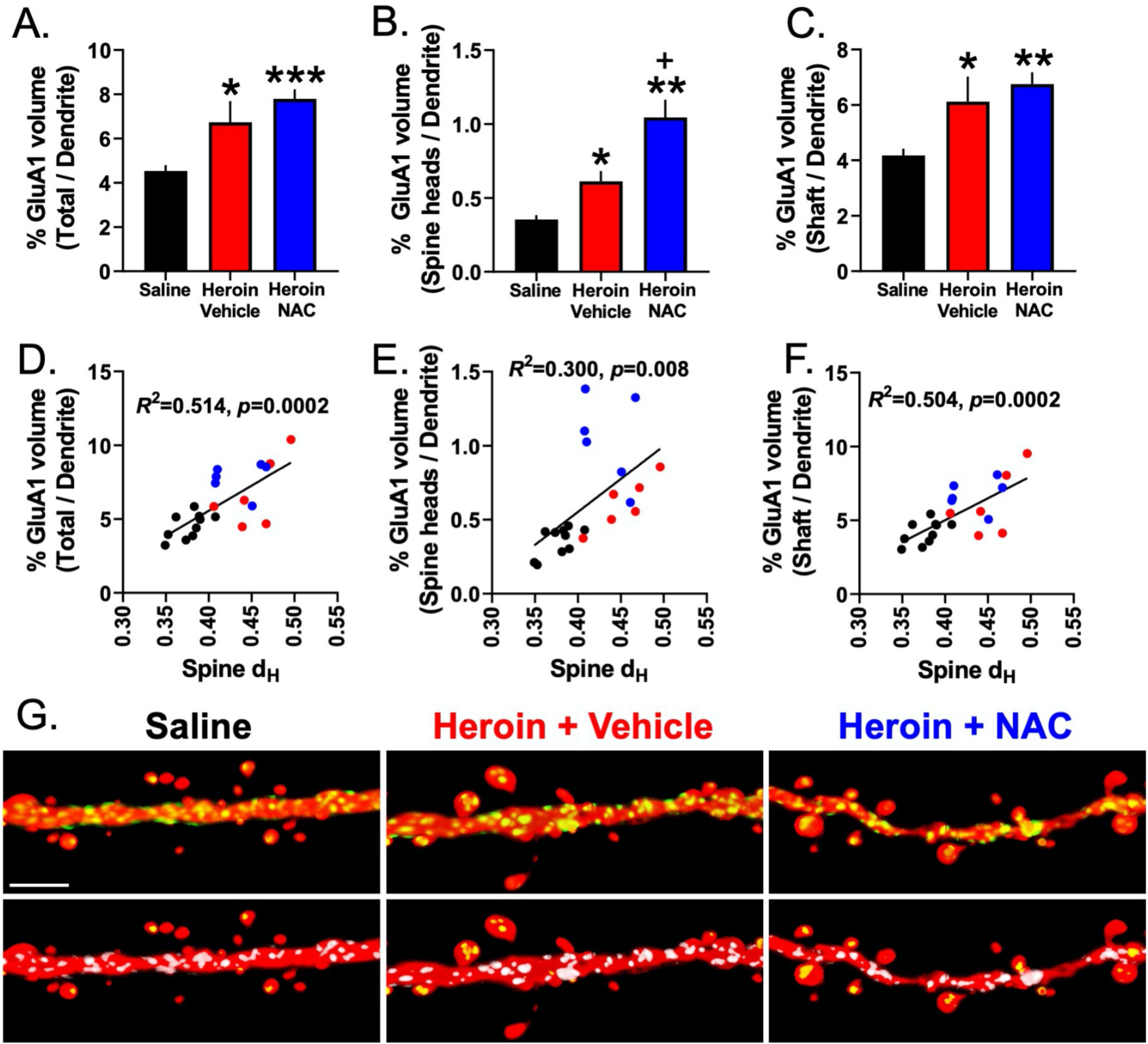
Heroin SA and extinction, regardless of NAC treatment, increased GluA1 immunoreactivity in dendritic shafts and spine heads of PrL-NAcore neurons. A. Heroin SA and extinction increased GluA1 immunoreactivity in dendritic spine heads and shaft, which was unaffected by repeated NAC treatment. B. Heroin SA and extinction increased GluA1 immunoreactivity in dendritic spine heads, which was further potentiated by repeated NAC treatment. C. Heroin SA and extinction increased GluA1 immunoreactivity in the dendritic shaft, which was unaffected by chronic NAC treatment. D-F. GluA1 immuonreactivity in the total dendrite (D), the dendritic spine heads only (E), as well as GluA1 immunoreactivity confined to the dendritic shaft (F), positively correlated with treatment-induced alterations in spine head diameter. G. Top panel – representative dendritic spine segments showing Flag (red) and GluA1 (green). For clarity, GluA1 is only shown in the volume occupied by the dendrite segment. Bottom panel – co-registry between GluA1 and Flag in the dendritic shaft (white) and spine heads (yellow). \**p*<0.05, \*\**p*<0.01, \*\*\**p*<0.001 compared to Saline; +*p*<0.05 compared to Heroin + Vehicle. Scale bar indicates 4 µm.

### Experiment 2 – Heroin SA and extinction increased GluA1 immunoreactivity in dendritic shafts and spine heads of PrL-NAcore neurons; and NAC potentiated spine head-specific GluA1

We previously showed that cocaine SA followed by one week of abstinence increases dendritic spine head diameter in apical tuft dendrites of PrL-NAcore neurons. We also demonstrated that cocaine abstinence duration-dependent structural plasticity occurs concomitantly with alterations in GluA1 immunoreactivity within the dendritic shaft and at dendritic spine heads [30]. Here, we investigated whether the adaptations in dendritic spine morphometrics following extinction of heroin SA (with or without repeated NAC treatment) were associated with altered GluA1 immunoreactivity in the discrete dendritic subcompartments of PrL-NAcore apical tuft dendrites described above. When analyzing total volume of GluA1 immunoreactivity in dendritic spine heads and shaft combined, relative to the total dendrite volume, a one-way ANOVA revealed a significant effect of treatment (F(2,19)=10.56, *p*=0.0008). Tukey’s multiple comparison test indicated that the Heroin + Vehicle (*p*=0.02) and Heroin + NAC (*p*=0.0009) groups displayed increased GluA1 immunoreactivity compared to saline controls **(Figure 5A)**. In the case of the Heroin + NAC, but not Heroin + Vehicle, group increases in GluA1 volume could occur in part due to increased overall GluA1 signal intensity, as the average GluA1 intensity (Spine heads + dendritic shaft) was significantly higher in this group (Main effect of treatment: (F(2,19)=4.27, *p*=0.029, Tukey’s post hoc comparing Heroin + NAC to Heroin + Vehicle – *p*=0.025, **Figure S3B)**.

Next, we focused on dendritic spine head-specific GluA1 immunoreactivity. Here the volume of signal in spine heads was expressed relative to the total dendrite volume, as a function of treatment. A Brown-Forsythe ANOVA revealed a significant effect of treatment (F(2,9.028)=20.22, *p*=0.0005). Dunnett’s T3 multiple comparison test revealed that the Heroin + Vehicle (*p*=0.03) and Heroin + NAC (*p*=0.004) groups showed increased spine head-specific GluA1 signal accumulation when compared to the Saline + Vehicle group **(Figure 5B)**. However, in this instance there was no difference between groups in the average GluA1 intensity in the co-registered volume **(Figure S3C)**.

Finally, we analyzed dendritic shaft-specific GluA1 immunoreactivity, relative to the total dendrite volume, as a function of treatment. A one-way ANOVA revealed a significant effect of treatment (F(2,19)=7.73, *p*=0.004). Tukey’s multiple comparison test indicated that the Heroin + Vehicle (*p*=0.033) and Heroin + NAC (*p*=0.005) groups showed increased dendritic shaft-specific GluA1 immunoreactivity compared to the Saline + Vehicle group **(Figure 5C)**. The Heroin + NAC group displayed increased GluA1 intensity in the dendritic shaft sub compartment compared to both Saline + Vehicle and Heroin + Vehicle groups **(Figure S3D)**. This is to be expected given the results discussed above as the shaft region constitutes a large portion of the total dendritic segment volume. Interestingly, correlation analyses revealed that % GluA1 volume in the total segment normalized to the total dendrite volume (*R*^*2*^=0.51, *p*=0.0002, **Figure 5D)**, % GluA1 volume in the spine heads normalized to the total dendrite volume (*R*^*2*^=0.300, *p*=0.008, **Figure 5E**), as well as % GluA1 volume in the shaft normalized to the total dendrite volume (*R*^*2*^=0.504, *p*=0.0002, **Figure 5F**) showed a positive correlation with changes in spine head diameter as a function of treatment. In contrast, the % GluA1 volume in the total dendrite, spine heads only, and shaft only, did not correlate with changes in dendritic spine density as a function of treatment **(Figure S4A-C)**. Representative dendritic spine segments with GluA1 immunoreactivity in the two sub compartments are shown in **Figure 5G**.

## Discussion

Our data demonstrate that heroin SA followed by extinction enhances the fundamental cytoskeletal cellular complexity of astrocytes in the PrL cortex, a finding that was accompanied by increased association of astrocytic processes with two distinct synaptic markers. Both of these effects, as well as heroin seeking during extinction, were normalized by repeated NAC treatment during the last 10 days of extinction training, confirming that NAC reduces responding during extinction as others have reported [19]. Further, these data implicate the heroin SA and extinction-mediated structural adaptations in astrocytes discussed above in the pathophysiology of heroin relapse vulnerability.

Apart from impacts on PrL cortical astrocytes, heroin SA and extinction also increased the average spine d_H_ on apical tuft dendrites of PrL-NAcore neurons, an effect that was accompanied by an overall decrease in dendritic spine density. Akin to its restorative effects on astrocytes, NAC treatment during extinction normalized heroin-induced decreases in dendritic spine density. However, NAC treatment *did not* impact elevations in dendritic spine d_H_. Further, NAC treatment actually potentiated the accumulation of GluA1 immunoreactivity in dendritic spine heads beyond levels observed after heroin SA and extinction. These results suggest that the ability of NAC to prevent heroin relapse in preclinical models may not only be due to its action in the NAcore but may also include actions that normalize structural plasticity in astrocytes, as well as neurons within the PrL cortex that project to the NAcore. While a causal role for these neuronal adaptations have yet to be established, NAC appears to function in the PrL cortex through mechanisms that extend beyond simple reversal of neuronal drug-induced structural plasticity.

### Repeated NAC treatment facilitated extinction training

Our data provide the first preclinical evidence for NAC acting to augment structural plasticity in a cortical region; we suspect that these adapatations play a role in NAC’s ability to prevent cue-induced reinstatement. Although NAC’s ability to promote abstinence has yet to be tested in the clinic for heroin-addicted patients, preclinical data indicate that daily NAC treatment during extinction reduces extinction responding as well as cue–and heroin– induced drug seeking for up to 40 days after the last injection [19]. Although rats did not undergo reinstatement in the current study, our data demonstrate that active lever presses over the last 3 days of extinction training are indeed reduced by repeated NAC treatment. Ultimately, NAC has shown promise preclinically in decreasing relapse for several drugs of abuse [14, 20, 22-24]. However, it is critical to note that NAC’s ability to suppress relapse clinically, at least in the case of cocaine, is minimal to date [36]. The general consensus is that NAC is effective at increasing abstinence duration in patients who are already abstinent, but fails to decrease relapse rates in active users [36-38]. Despite its modest clinical efficacy, its well-described mechanism of action and ability to target drug-induced adaptations in non-neuronal cells substantiates its role as a powerful research tool for understanding astrocytic dysfunction in addiction. Regarding treatment groups, in the current study we chose to omit Yoked-Saline animals injected with NAC for several reasons. Most importantly, NAC has been shown to have no effect on NAcore glutamate concentrations or cystine uptake through X_c_^-^ in control animals [39]. Moreover, NAC treatment normalized increased glutamate concentrations in the anterior cingulate of cocaine-addicted patients but had no effect in healthy controls [40].

### Repeated NAC treatment during extinction reversed increases in astrocyte complexity and association of PAPs with synapses in PrL cortex induced by heroin SA and extinction

Here we provide evidence that extinction from heroin SA increased GFAP cytoskeletal complexity without altering the number of GFAP+ cells in the PrL cortex. We conclude that this is not due a transition of astrocytes to a reactive state as there was no enhancement of average GFAP filament diameter across groups, a phenomenon that serves as a canonical index of reactive astrocytes [41]. Given the profound impacts observed in PrL cortical astrocytes, the signal transduction required for this event is an immediately apparent question. It is possible that this effect may be due to heroin’s direct impact on opioid receptors expressed on astrocytes. mRNA for all three opioid receptor subtypes are present in astroglia and PrL cortical astrocytes in particular express an abundance of mu opioid receptor mRNA [42], the primary receptor engaged by morphine [43]. Further, incubation of cortical astrocyte cultures with morphine enhances intracellular Ca^2+^ signaling [44], resulting in increased area occupied by astrocytes [45]. In addition, astrocytic mu opioid receptor expression has been directly linked to the rewarding properties of drugs of abuse whereby activation of mu opioid receptors on hippocampal astrocytes induces conditioned place preference [46]. Thus, it is likely that one component of the observed increase in GFAP complexity in the PrL cortex of heroin-exposed rats is due to chronic tone on astrocytic mu opioid receptors. In support, a study performed in the early 1990’s showed that cultured astrocytes treated with morphine increase their branching complexity; yet morphine decreased astrocyte proliferation [27]. Alternatively, chronic injections of morphine in rats have been shown to increase GFAP protein expression in the frontal cortex [26], yet this study did not have a prolonged withdrawal period. While we did not causally link increased GFAP complexity to relapse behavior, repeated NAC treatment prevented the action of heroin at cortical astrocyte GFAP arbors and is known to reverse drug-induced deficits in astrocyte function linked to relapse propensity [39, 47]. Whether normalized cortical GFAP complexity is required for NAC’s ability to suppress relapse remains to be determined experimentally.

Extinction from cocaine [13, 34], heroin [12], or methamphetamine [14] SA leads to decreased association of PAPs with synaptic markers in the NAcore. This finding can be interpreted as a withdrawal of PAPs, which are highly enriched in GLT-1 [48], from synapses [49, 50]. This connection between astrocyte structure and function is readily made as drug-induced retraction of glial processes in the NAcore is often paired with decreased GLT-1 expression and reduced glutamate clearance [51]. We show here that extinction from heroin SA leads to increased association of PAPs with synaptic markers in the PrL cortex, an effect that was specific to heroin and was not observed following extinction of cocaine SA [34]. While further investigation would be required to determine if heroin SA and extinction impacts cortical glutamate clearance, we would not predict reduced glutamate clearance, as astrocyte retraction from synapses was not observed. The GFAP cytoskeleton represents a long standing canonical maker for astrocytes and has been often utilized as a means to interrogate astrocyte structure [52, 53] and activation states [54]. However, it of course does not accurately recapitulate the membranous extremities of astrocytes [55] that extend beyond the GFAP skeleton. Accordingly, alterations in the cytoskeletal properties of astrocytes at the level of GFAP arbors do not necessarily mandate that adaptations in astrocyte-synaptic interaction will be observed. As an example, methamphetamine SA and extinction decreased PAP-synapse association, but not GFAP cytoskeletal complexity, in the NAcore [14]. Here we do observe that heroin-induced increases in GFAP complexity were accompanied by ∼75% increase in the association of PAPs with synaptic markers in the PrL cortex following extinction from heroin SA. Interestingly, both adaptations in GFAP structure and astrocyte-synaptic interaction were normalized by repeated NAC treatment. One potential alternative explanation for our observations is that there was increased synaptogenesis in the PrL cortex after extinction of heroin SA, with no alteration in the motility of astrocytes. However, this is unlikely given that we did not observe any difference between groups in the number of GluA2 or Synapsin I puncta.

It has been estimated that a single astrocyte makes contacts with ∼100 dendrites and ∼100,000 synapses in the rodent brain [56]. Our astrocyte-synaptic co-registration data could be interpreted either as each astrocyte contacting a greater number of synapses in the PrL cortex after extinction or that each astrocyte may be embracing individual synapses to a greater extent. Additionally, some combination of these scenarios may also be occurring. Each of these potential outcomes would produce an overall increase in co-registry of astrocyte membrane and synaptic marker signals. Astrocyte ensheathment of synapses shapes excitatory transmission by establishing the three-dimensional space in which neurotransmission occurs in. More specifically, astrocyte ensheathment of synapses is a dynamic process that regulates the profiles of synaptic and non-synaptic receptor activation, allowing for astroglial tuning of synaptic communication on a synapse-by-synapse basis. This modality of regulation has been directly explored by studies that examine how the degree to which astrocytes associate, and physically interact, with synapses is shaped by neuronal activity [57, 58]. As an example, whisker stimulation has been shown to increase GLT-1 expression in the barrel cortex, which is associated with increased ensheathment of PAPs at enlarged dendritic spine heads [59]. Additionally, cue-induced reinstatement of heroin seeking has been associated with dynamic changes in PAP-synapse association in the NAcore during the course of the reinstatement session that was directly linked to relapse behavior [12]. Together these data demonstrate that astrocytes physically respond to changes in neurotransmission with physiologically and behaviorally relevant consequences. Moreover, the fidelity of synaptic transmission is tightly controlled by PAP insulation of synapses; too little association is linked to glutamate overflow [12-14] whereas too much insulation can actually impede AMPA receptor-mediated excitatory transmission through enhanced glutamate uptake [60], a means of astroglial regulation of synaptic function initially postulated to occur during sleep by Cajal almost 150 years ago [61]. Further experimentation will be needed to determine where astrocytic synapse ensheathment in the PrL cortex from heroin-experienced animals lies on this functional continuum. Whether PAP association with synapses has reached a “ceiling” and is impeding AMPA receptor transmission or whether the increased association leads to a suppression of volume transmission and a solidification of discrete excitatory inputs in a supraphysiological manner is yet to be determined.

### Extinction from heroin SA altered dendritic spine morphological profiles and GluA1 immunoreactivity in discrete dendritic subcompartments of prelimbic cortical neurons projecting to the nucleus accumbens core

Using similar viral vector-based pathway-specific labeling of dendritic spines, we previously found that cocaine SA followed by one week of abstinence decreased the density of apical tuft dendritic spines of PrL-NAcore neurons, yet the remaining spines were enlarged and concomitantly displayed increased GluA1 and GluA2 immunoreactivity in the spine heads [30]. A previous study found that heroin SA and one month of abstinence decreased the density of dendritic spines of PrL cortical layer V pyramidal neurons, with no reported alterations in morphometric properties of the spines themselves [35]. To accompany the astrocyte-synapse interaction data discussed above, we investigated potential alterations in dendritic spine morphological profiles in PrL cortical neurons projecting to the NAcore. Akin to one week of abstinence from cocaine SA, we discovered that extinction from heroin SA decreased the density of dendritic spines along dendrites at the apical tuft of PrL-NAcore neurons, yet these neurons showed an increase in the average spine head size as well as elevated GluA1 co-registry in the dendritic spine heads. These data are largely consistent with studies demonstrating repeated treatment of cultured cortical neurons with morphine leads to destabilization of pre-existing spines [62]. In our data set, heroin-mediated decreases in dendritic spine density were prevented by repeated NAC treatment; yet the heroin-evoked increases in spine head diameter were not. In fact, to our surprise, we found that NAC treated animals showed a potentiation of GluA1 immunoreactivity in dendritic spine heads relative to Heroin + Vehicle animals. Generally, an increase in the number of spines, the average spine head size, as well as AMPA receptor occupancy in spine heads, is interpreted as evidence for synaptic potentiation [63-66]. Thus, this finding is initially counter-intuitive given that NAC normalized the decrease in density, but NAC-treated animals retained enlarged dendritic spines following heroin SA and extinction, suggesting that NAC is driving synaptic potentiation in the PrL-NAcore circuit. However, it should be noted that the apical tuft is a site of convergence of glutamatergic inputs from midline thalamic nuclei as well as dopamine afferents from the VTA[67-69]. Thus, it remains possible that heroin-induced adaptations in spine morphology and AMPA receptor accumulation at this particular part of the dendritic arbor might not translate into whole cell alterations in physiological indices of potentiation. Rather, this may reflect potentiation of discrete inputs onto PrL-NAcore neurons which may or not may be counteracted by alterations in synaptic strength of alternative inputs at other parts of the dendritic arbor. Future studies should examine whether heroin SA and extinction alter the electrophysiological properties of PrL-NAcore neurons and whether repeated NAC treatment during extinction has an impact on such alterations. Regardless, given that NAC is known to reverse drug-induced deficits in astrocyte function [18], we suspect that NAC’s ability to restore dendritic spine density following heroin SA and extinction is mediated by augmented astrocyte function, perhaps through altering the release patterns of astrocyte-derived neuromodulators that regulate synaptogenesis.

It is well-documented that astrocytes release a variety of neurotransmitters, neuromodulators, and growth factors that can have a profound impact on neuronal plasticity, (for review see [70, 71]). As an example, astroglial release of thrombospondin-1 (TSP-1) activates α2δ-1 receptors expressed on neurons to induce spinogenesis and excitatory synapse formation [72]. Dysfunction of this system has been linked to reduced dendritic spine density in down syndrome models [73], as well as elevated dendritic spine density in the NAc of animals receiving chronic cocaine injections[74]. It is possible that the heroin-induced decrease in spine density could be linked to dysfunctional gliotransmission following extinction, and that NAC’s ability to drive spinogenesis could be linked to normalization of such dysfunction. For example, repeated morphine administration in mice downregulates TSP expression in the cortex 24 hours after the final injection, as well as in cultured astrocytes following repeated morphine treatment [75]. One mechanism by which heroin may impact TSP expression is through increased oxidative stress, as evidenced by the induction of oxidative stress in humans following chronic heroin exposure [76] and also in cultured cortical neurons [77, 78]. Moreover, cultured primary astrocytes reduce TSP-1 gene expression under increased oxidative stress conditions, which is prevented by NAC [79]. Given that one well-documented mechanism by which NAC counteracts oxidative stress is increased glutathione synthesis [80], it is possible that heroin-induced adaptations in TSP-1 expression due to increased oxidative stress, and normalization of which by NAC, may be linked to the altered dendritic spine morphometric features observed herein. Overall, it is likely that NAC’s ability to normalize heroin-induced adaptations in astrocyte-synapse proximity and GFAP cytoskeletal architecture has a direct influence on structural plasticity in PrL-NAcore neurons, perhaps through indirect or direct mechanisms at neurons. Thus, the degree to which PAPs associate with dendritic spines of PrL-NAcore neurons, whether this is altered following extinction of heroin SA, and whether NAC has an impact are areas of future investigation.

### Methodological considerations

Canonically, GFAP is thought to only be present in the major branches and endfeet of astrocytes and not to be present in all non-reactive astrocytes in adult healthy CNS tissue [81]. However, our data are potentially at odds with this view as we found consistent fields of astrocytes in all three experimental groups and did not observe gaps that could potentially represent GFAP-negative astroglia. As described above, GFAP does not extend into the finest membranous processes, which are the primary domains of astrocyte-neuron interactions; the motility of which are regulated by actin cycling [82]. The ability to use membrane-targeted expression of GFP driven by astrocyte-specific promoters in conjunction with immunohistochemical detection of synaptic proteins has gained traction as an assay to determine the proximity of PAPs to synapses (see [50] and [49] for reviews). In this study, we used two different synaptic markers; Synapsin I and GluA2 which label generic pre-synaptic terminals (Synapsin I) and excitatory post-synaptic AMPA receptors (GluA2), respectively. However, it is important to note that with an approximate minimum resolution of 150 nm, we are not attempting to distinguish between pre-and post-synaptic elements; separated by an approximate distance of 10 nm [83]. We are merely using IHC detection of these proteins as a means to label synapses. However, the fact that both antibodies yielded increased co-registration of synapses with astrocytic membranes in heroin animals can be viewed as a means to exclude epiphenomenological or potentially off-target effects from each individual antibody.

## Conclusions

In conclusion, our data indicate that heroin SA and extinction elicit hypertrophic complexity and synapse association of astroglia in the PrL cortex that are associated with altered dendritic spine morphological features in PrL-NAcore neurons. Our data fill a much-needed gap in the field by focusing on the upstream portion of the canonical relapse circuit to accompany what has been observed in the NAcore. Moreover, these findings point to an additional node of this circuitry whereby NAC can modify heroin-induced adaptations in the PrL cortex in addition to its well-defined modification of NAcore synaptic pathology. Ultimately, astroglia and neuronal adaptations in the PrL cortex may be a target for informing future drug discovery for relapse prevention pharmacotherapies.

## Acknowledgements

This was work was supported by R00 DA040004 (MDS), F32 DA050427 (BMS), T32 DA007288 (JFM, BMS), P50 DA046373 (JFM, MDS). This publication was supported by the South Carolina Clinical & Translational Research (SCTR) Institute, with an academic home at the Medical University of South Carolina CTSA, NIH/NCATS Grant Number UL1 TR001450. The contents are solely the responsibility of the authors and do not necessarily represent the official views of the NIH or NCATS. The authors thank Dr. Giuseppe Giannotti for excellent technical assistance.

## Supplemental Data

**Figure S1.**
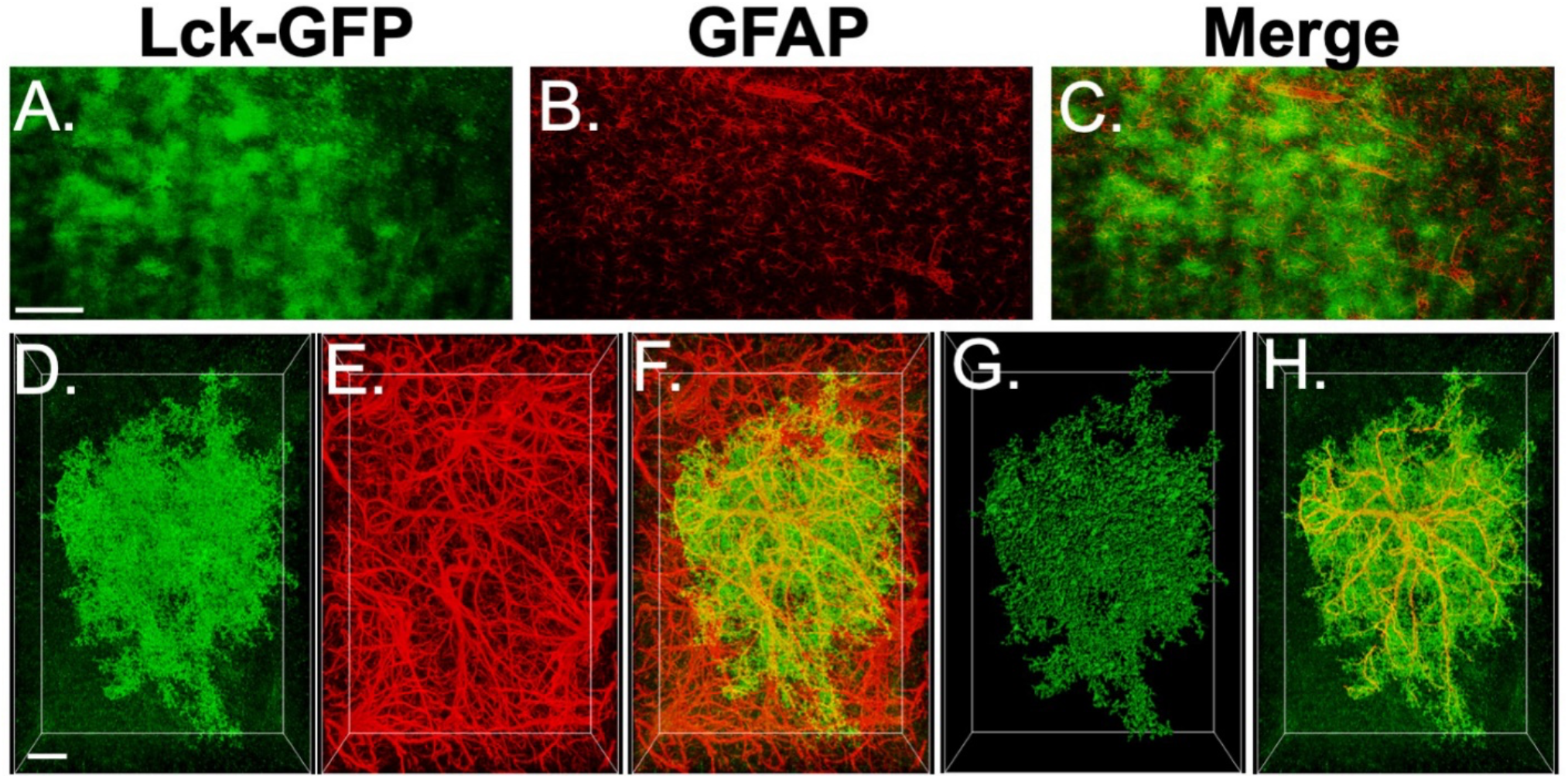
AAV5-GfaABC1D-Lck-GFP is specific to GFAP+ astrocytes. A-C. Low magnification view of AAV5-GfaABC1D-Lck-GFP, GFAP, and the merge of the two signals. D-F. Raw deconvolved GFP, GFAP immunohistochemical detection, and the merge of the two signals. G. 3D space filling model to isolate the GFP+ astrocyte from background. H. GFAP signal confined to the isolated GFP+ astrocyte. Scale bars indicate either 150 µm (A-C) or 10 µm (D-H).

**Figure S2.**
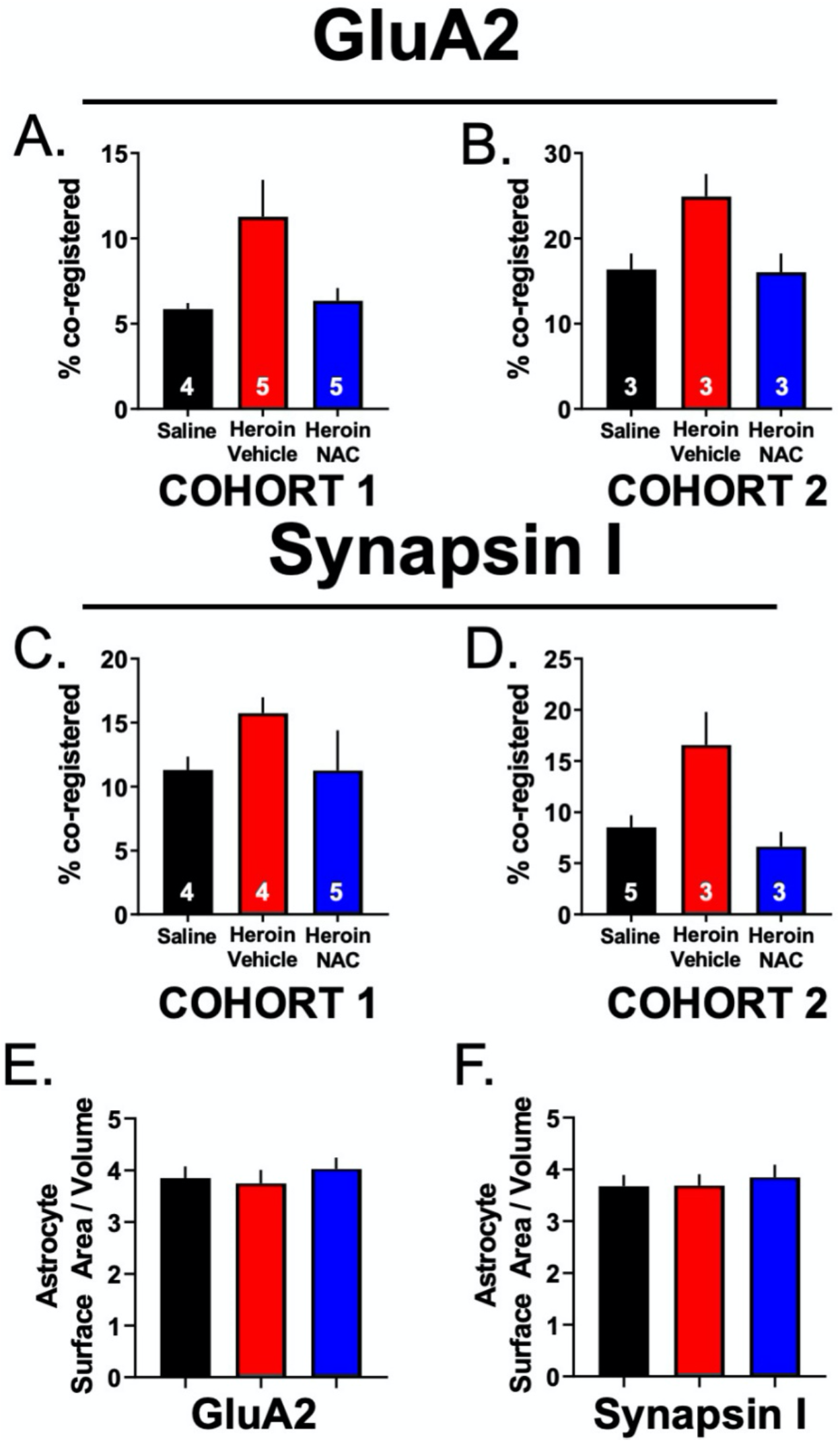
Between cohort effects on GluA2 and Synapsin I co-registry with Lck-GFP. A-B. Cohort 1 (A) and cohort 2 (B) showed the same effect of percent co-registry for GluA2. C-D. Cohort 1 (C) and cohort 2 (D) showed the same effect of percent co-registry for Synapsin I. E-F. 3D rendered Lck-GFP-expressing astrocyte showed no difference between groups in the surface area to volume ratio for GluA2 (E) or Synapsin I (F) analyses.

**Figure S3.**
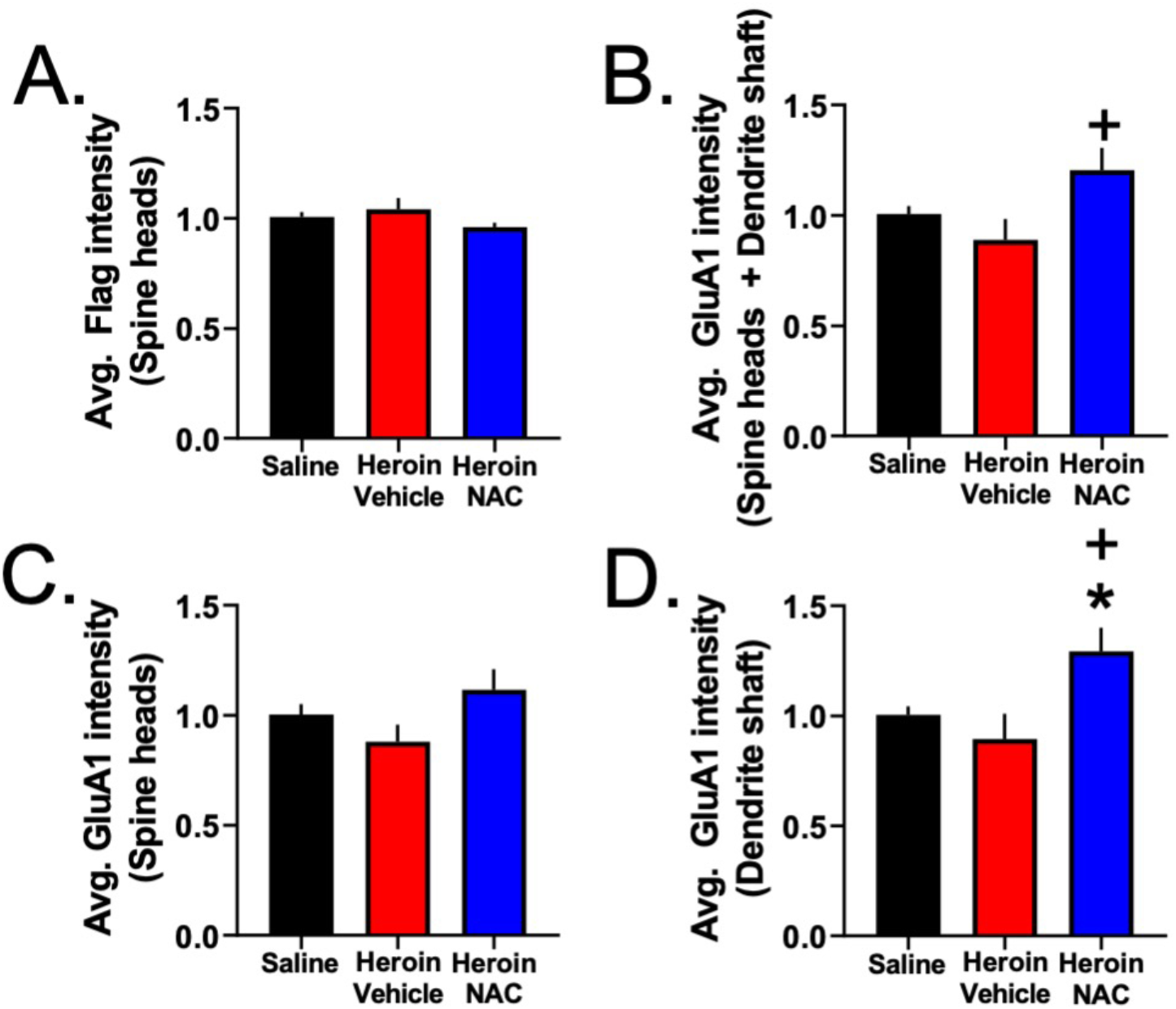
Signal intensity data pertaining to Experiment 2. A. There was no difference between groups in the average Flag intensity in dendritic spine heads. B. Heroin + NAC group showed increased GluA1 intensity in the co-registered volume within spine heads and shafts (combined) compared to Heroin + Vehicle, but not saline. C. There was no difference between groups in the average GluA1 intensity in the volume of co-registered dendritic spine heads. D. Heroin + NAC group showed increased GluA1 intensity in the co-registered volume within the dendritic shaft compared to Saline and Heroin + Vehicle. \**p*<0.05 compared to Saline; +*p*<0.05 compared to Heroin + Vehicle. Data is expressed as a fold change from control.

**Figure S4.**
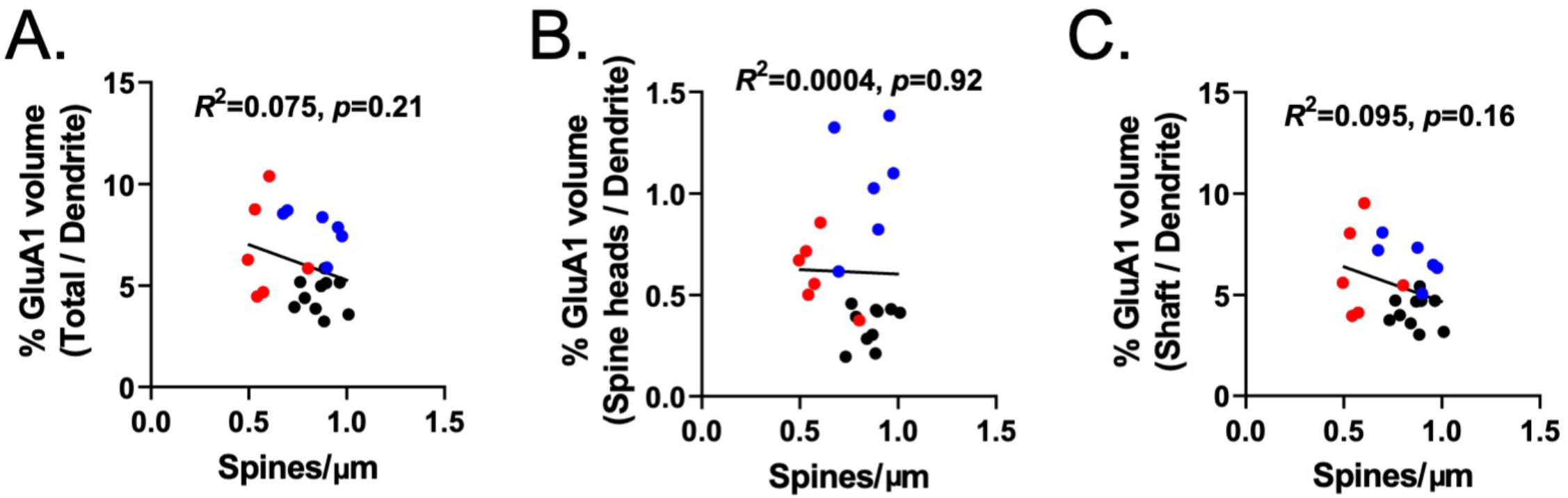
There was no correlation between the percent GluA1 volume in subcompartments normalized to the total dendrite volume and spine density. A-C. % GluA1 volume in the co-registered volume within the total dendrite (A), spine heads only (B), or shaft only (C) was not predictive of changes in spine density as a function of treatment.

